# Cortical response to balance perturbation is more sensitive in modern dancers than nondancers during biomechanically similar balance recovery

**DOI:** 10.1101/2025.11.14.688528

**Authors:** Kennedy G. Kerr, Scott E. Boebinger, Jasmine L. Mirdamadi, Janna Protzak, Michael R. Borich, Lena H. Ting

## Abstract

Motor skill expertise can facilitate more automatic movement, engaging less cortical activity while producing appropriate motor output. Accordingly, cortical-evoked N1 responses to balance perturbation, assessed using electroencephalography (EEG), are smaller in young and older adults with better balance. These responses may thus reflect individual balance challenge versus functional, or objective, task difficulty. However, the effect of balance expertise on cortical responses to balance perturbation has not been studied. We hypothesized that balance ability gained though long-term training facilitates more automatic balance control. Using professional modern dancers as balance experts, we compared cortical-evoked responses and biomechanics of the balance-correcting response between modern dancers and nondancers. We predicted that modern dancers would have smaller cortical-evoked responses and better balance recovery at equivalent levels of balance challenge. Support-surface perturbations were normalized to individual challenge levels by delivering perturbations scaled to 60% and 140% of each individual’s step threshold. In contrast to our prediction, dancers exhibited larger N1 responses compared to nondancers while demonstrating similar biomechanical responses. Our results suggest dancers have greater cortical sensitivity to balance perturbations than nondancers. Further, dancer N1 responses modulated across perturbation magnitudes according to differences in objective task difficulty. In contrast, nondancer N1 responses modulated as a function of individual challenge level. Our findings suggest dance training increases sensitivity of the initial, cortical N1 response to balance perturbation, supporting postural alignment to an objective reference. The N1 response may reflect differences in balance-error processing that are altered with specific long-term training and may have implications for rehabilitation.

**NEW & NOTEWORTHY:** Modern dancers show larger cortical responses to balance perturbations than nondancers, suggesting a greater sensitivity to perturbations. These results contrast with evidence of larger cortical-evoked responses in young adults with poorer balance, consistent with the cortical N1 response being a balance error assessment signal. Whereas nondancers scaled cortical responses by individual differences in N1 amplitude, dancers’ cortical responses were scaled to objective differences in perturbation magnitude, suggesting increased postural awareness due to training.

## INTRODUCTION

Expertise in motor skills is associated with automatic motor control, requiring less cortical activity to produce the appropriate motor output (1–4). For example, cortical activity is reduced in musicians performing skilled voluntary tasks (5, 6). However, it is unclear whether this phenomenon applies to whole-body movement. Professional modern dancers are a cohort of individuals trained in whole-body motor control (7, 8). Although dance-based rehabilitation is currently used for balance and motor impairment (9–13), few studies have investigated cortical activity and biomechanics concurrently during whole-body movement in dancers (14, 15). Previous investigations into dancer motor control were limited by exploring either cortical activity of dancers during action observation (16, 17) or at rest (18) or biomechanics of dancers during walking (19), but not both together. Studying the effects of balance-improving motor skill on whole body neurophysiology and biomechanics could provide novel insight into the potential role of balance training in rehabilitation.

Balance control is typically thought to be an automatic task engaging primarily subcortical control (20), but measuring cortical activity during balance can provide insight into hierarchical sensorimotor control loops also engaged (21). In older adults, balance performance decreases when a concurrent cognitive task is performed as assessed via center of pressure (CoP) metrics during quiet stance (22, 23). These results suggest finite cortical resources can become engaged during balance and may not be able to compensate when a concurrent task engages the same resources (24). These indirect measures of cortical engagement have been corroborated by electroencephalography (EEG) studies measuring increased cortical activity during reactive balance between older adults with clinical balance impairment compared to age-matched controls (25–27). In young and older adults, increased cortical activity measured with EEG or functional near-infrared spectroscopy is observed within individuals as balance challenge increases (28–32). Many studies have analyzed the cortical N1, the first negative event related potential (ERP) following balance perturbation occurring ∼150ms after perturbation and localized to the supplementary motor area (33, 34). The N1 is thought to be a balance error assessment signal (35, 36), increasing with perturbation difficulty (29, 33, 35, 37–40) or perceived threat (41, 42). There is also conflicting evidence for whether the N1 Is associated with the allocation of attentional resources toward balance (43).

Evoked N1 potentials during balance perturbation have been associated with differences in balance ability in young and older adult cohorts, demonstrating the need to account for balance ability in determining perturbation challenge. N1 responses to height-scaled perturbations are larger in individuals with lower balance ability and older adults with lower self-reported balance confidence (33, 38). This shows that height normalized perturbations control for biomechanical constraints of an individual but not the effect of balance ability on perturbation difficulty, or functional task difficulty (44). Here we use the term “balance challenge” to refer to the combination of the effects of height and balance ability on perturbation difficulty. One way to identify individual-specific balance challenge posed by a task is by determining step threshold, the maximum perturbation magnitude an individual can withstand without necessitating a step to recover balance (22). N1 amplitude is larger during stepping responses than non-step responses at the same perturbation magnitude (29, 45), but N1 potentials have not been compared across individual balance challenge. To compare balance error assessment across levels of motor skill training, we delivered perturbations at equal levels of balance challenge across participants by normalizing perturbations to step threshold. Step threshold scaling allows for between-subjects comparisons at the same challenge level, and within-subjects comparisons across challenge levels.

We hypothesized that balance expertise leads to more automatic balance control requiring less cortical engagement. We tested professional modern dancers as a cohort of individuals with balance expertise and evaluated their cortical N1 responses to balance perturbations. Perturbations were normalized to individual step threshold to ensure a similar level of balance challenge across participants and isolate the effect of expertise on balance error assessment between professional modern dancers and nondancers. We predicted balance perturbations would evoke smaller cortical N1 responses in experts compared to novices when they were similarly challenged. We then tested whether the magnitude of the N1 responses scaled with balance ability as assessed by a narrowing beam walking task (46). Finally, we tested whether smaller N1 responses were associated with faster biomechanical responses to perturbations as measured by CoP rate of rise (RoR) (47, 48).

## MATERIALS AND METHODS

### Participants

Data was collected from 10 professional modern dancers (20-32 years old, 5F) and 14 novice healthy young adults (21-29 years old, 7F) recruited from Emory University and the surrounding Atlanta community. All participants had no history of neurological, musculoskeletal, and/or visual impairments. Inclusion criteria for dancers was at least 10 years of pre-professional and/or professional training. All participants provided written informed consent in line with the Emory Institutional Review Board before beginning the study. Of the collected data, 2 male dancers were excluded for poor EEG quality relating to hardware issues, and 2 female nondancers were moved to the dancer group upon learning these individuals had received at least 10 years of dance training, though not at the professional level. Thus, data from 10 dancers (7F) and 12 nondancers (5F) is analyzed here.

### Step threshold

To ensure subsequent support surface translation perturbations could be scaled to challenge level across participants, we first determined individual step threshold, or the perturbation magnitude that induces a step to recover balance 50% of the time (49). Participants were given 30 forward support surface translation perturbations (Fig. 1) (AMTI force plate instrumented custom platform, Factory Automation Systems, Atlanta GA) at unpredictable timing and magnitude along with randomly delivered backward perturbations to limit habituation. Perturbation magnitude adaptively iterated above and below the current predicted step threshold using a previously published method (49). Participants were instructed to recover balance without taking a step with eyes open and arms crossed across the chest. Participants wore a harness attached to the ceiling and were barefoot for all perturbations. Step threshold magnitude was identified by the end of the 30 trials for all participants.

**Figure 1.**
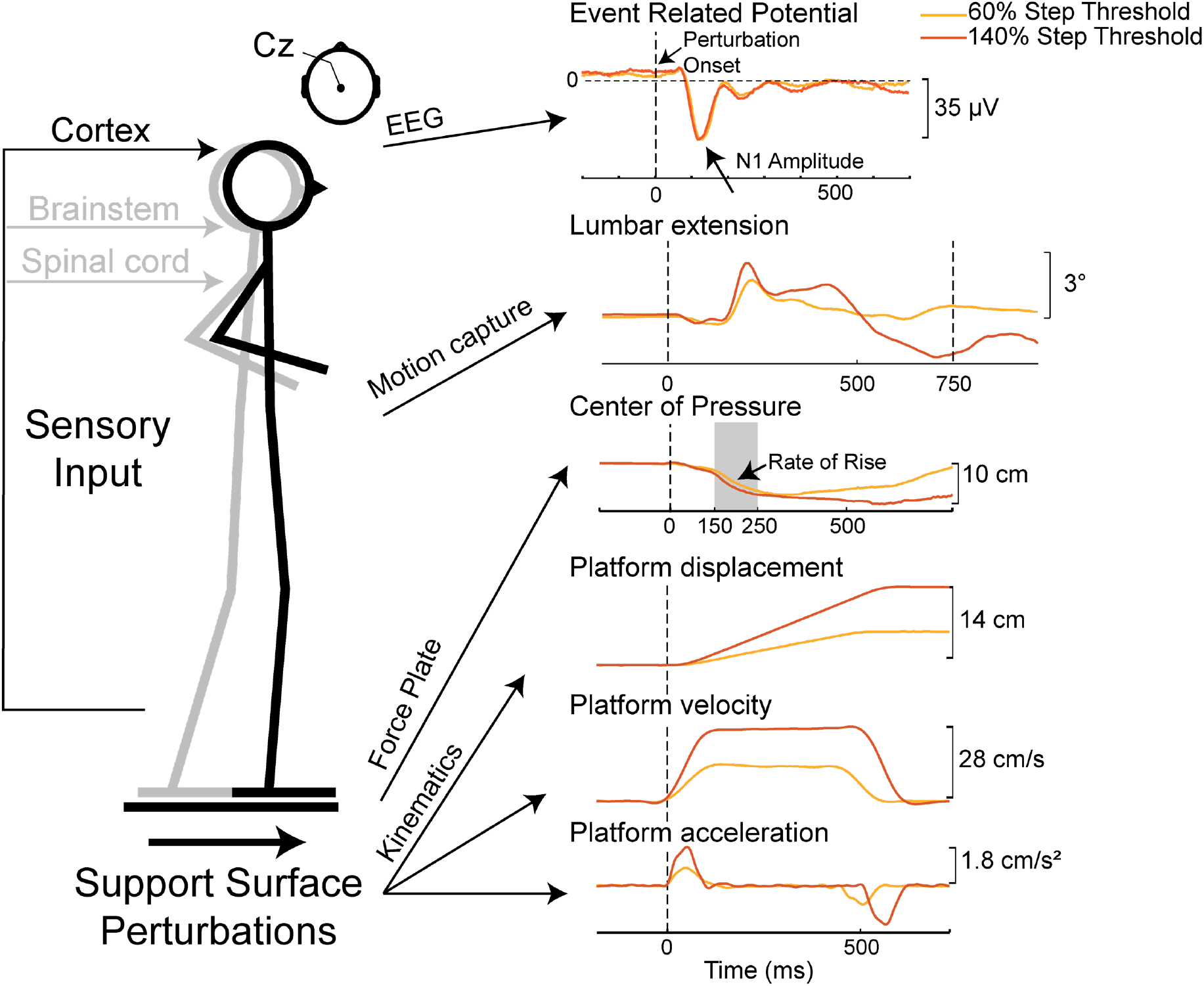
Experimental paradigm. Support-surface perturbations were given at two magnitudes scaled to individual step threshold. Surface EEG recorded evoked cortical responses throughout balance recovery. Lumbar extension angle, where positive values are extension and negative values are flexion, was calculated with inverse kinematics using 3D motion capture data. Center of pressure (CoP) displacement was calculated from force plates embedded in the platform, under each foot. CoP displacement 150-250 ms post-perturbation is used to calculate CoP rate of rise (RoR), or the efficiency of active balance-correcting CoP movement.

### Perturbation series

The step threshold magnitude determined in the first block of perturbations was used to scale the subsequent 72 perturbations to either 60% of the step threshold, ensuring a step is not needed to recover balance, or 140% of the step threshold, ensuring that a step is required to recover balance (Fig. 1). Participants were instructed not to take a step to recover balance unless necessary to prevent a fall. This created two challenge levels across participants: “no step” perturbations at 60% step threshold and “unplanned step” perturbations at 140% step threshold. 60% and 140% step threshold perturbations were delivered randomly and at unpredictable timing, along with small backward “catch” trials to limit habituation.

### Beam-walking task

To quantify balance ability, participants completed a challenging beam walking task (19, 46). Wearing standardized shoes and with their arms crossed, participants attempted to walk all the way across a narrowing balance beam (9.14 m long, 2.56 cm tall, segments narrowing from 18.6 cm to 2 cm wide every 1.83 m). Each of 6 trials ended when the participant reached the end, stepped off the beam, or uncrossed their arms. Beam walking was performed at self-selected speed, with the only objective of the participant being to reach the end of the beam. Balance ability was then quantified as “beam score”, or the mean normalized distance traveled across 6 trials (0 to 1). Participants were not instrumented with EEG during this task.

### Center of pressure (CoP)

6-axis forces under each foot were collected with 2 AMTI OR6-6 force plates within the perturbation platform at 100 Hz. We described onset of the balance correcting response to perturbations as center of pressure (CoP) rate of rise (RoR). This is the slope of the linear portion of CoP displacement over time between 150 and 250 ms after perturbation onset (Fig. 1). During this time bin, backward CoP movement is caused by active contributions to balance, counteracting passive forward CoP movement from the perturbation itself (47, 48). This time bin also occurs before foot-off of balance correcting steps that occurred in response to 140% step threshold perturbations. Faster CoP RoR represents faster response times and a more effective balance correcting response (48).

### Lumbar extension angle

Body motion was recorded at 100 Hz from a 33-marker set modified from the Vicon Dynamic Plug-In Gait model. Kinematics were calculated with marker data using the OpenSim Inverse Kinematics toolbox (50). Lumbar extension angle was baseline subtracted and quantified as the angle 750 ms after perturbation onset, or 250 ms after the end of the 500 ms perturbation (33).

### EEG recording and analyses

Cortical activity was recorded continuously throughout standing balance perturbations with a 64-channel active electrode cap and ActiChamp amplifier (Brain Products GmbH, Munich, Germany) according to the 10% system (51). Fz was the reference electrode. EEG data were processed using the Matlab EEGLAB toolbox (52) and custom Matlab scripts informed by steps in Makoto’s preprocessing pipeline (53). This preprocessing pipeline was described previously (54). Data were down sampled to 500 Hz and high-pass filtered with a 1 Hz cutoff. Channels were removed with the clean_rawdata plugin (4.4 ± 2.9 channels removed) if a channel was flat for more than 5 seconds, had a high-frequency noise standard deviation of more than 4, or had less than a 0.6 correlation with nearby channels (55). All removed channels were interpolated using a spherical function. Then the average reference was computed, 60 Hz line noise was removed with the Zapline plus plugin (56), and data were epoched 5 seconds before to 10 seconds after each perturbation and decomposed into maximally independent components using adaptive mixture independent component analysis (AMICA). Independent components were classified as brain sources with the ICLabel plugin and kept if they were classified ‘Brain’ with above 70% confidence and confirmed by visual inspection. Dipole locations were then estimated for these ICs using an MNI template with the DIPFIT plugin (57). ICs were further rejected if they were located outside of the brain or had a residual variance > 15% when autofitting dipoles (58). Channel-level data were reconstructed with the remaining neural components (7.0 ± 1.8 ICs back projected). Analyses focused on the Cz electrode. After baseline subtracting mean activity from 5 seconds to 1 second before the perturbation from the event related potential (ERP), N1 amplitude was defined as the minimum value of the trial-averaged ERP between 100 and 300 ms after the perturbation for each perturbation condition (Fig. 1) (40). We also analyzed the modulation of N1 amplitude between perturbation conditions (140% ST – 60% ST), where larger values indicate a greater change (ΔN1) between conditions for each individual.

### Statistical comparison across cohorts and perturbation magnitude

Two-sample t-tests were used to determine cohort-level differences in beam score, step threshold, and participant height. A repeated measures ANOVA was used to quantify differences in N1 amplitude by perturbation magnitude, cohort, and the effect of cohort on the relationship between N1 amplitude and perturbation magnitude. A similar repeated measures ANOVA was used to quantify differences in CoP RoR and lumbar extension at 750ms post-perturbation by perturbation magnitude, cohort, and the effect of cohort on any relationship between CoP RoR or lumbar extension and perturbation magnitude. Linear mixed effects models on single trial outcome measures and estimated marginal means were used for post-hoc comparisons between N1 amplitudes, CoP RoR, and lumbar extension between perturbation magnitudes and cohort.

### Statistical comparison across balance ability

A multivariate linear regression was performed to investigate the relationship between N1 amplitude and beam score. For this analysis, beam score was separated into quartiles of balance ability as previous work has shown different associations between balance ability quartiles (33). Beam quartiles and cohort were fixed effects and participant was a random effect. Estimated marginal means were used in post-hoc analysis of the relationship between N1 amplitude and perturbation condition within each quartile of balance ability. Linear models were also used to compare N1 amplitude and beam score using the average N1 amplitude for all perturbations together. These analyses quantified relationships between N1 amplitude with beam score in the dancer and nondancer cohorts independently, as well as with all participants taken together. Linear models were also used to determine relationships between the change in N1 amplitude from the 140% step threshold to 60% step threshold conditions to step threshold identified during the step threshold perturbation series and N1 magnitude in the non-step, 60% step threshold perturbation condition. All metrics are reported as the mean ± standard deviation unless otherwise noted.

## RESULTS

Dancers had higher balance ability than nondancers. Dancers performed better on the narrowing beam walking task (beam score: dancers=0.71 ± 0.10, nondancers=0.63 ± 0.07, t=-2.7, p=0.01). Step thresholds and height were similar between dancers and nondancers (step threshold: dancers=14.6 ± 2.0 cm, nondancers=13.5 ± 3.5 cm, t=-1.3, p=0.2; height: dancers=167.9 ± 7.1 cm, nondancers=171.7 ± 11.8 cm, t=1.3, p=0.2).

N1 amplitudes were larger in dancers than nondancers and scaled with changes in perturbation magnitude in both cohorts (Fig. 2A/B). In an exemplar dancer and nondancer who had the same step threshold and thus experienced the same perturbation magnitudes, the dancer exhibited larger N1 amplitudes in response to both conditions (Fig. 2A), consistent with N1 amplitude differences between cohorts (dancers= 47.5 ± 23.7 μV, nondancers=37.9 ± 21.1 μV, p=0.048). In these exemplars, the nondancer appears not to scale N1 responses between perturbation conditions (Fig. 2A). However, across cohorts, N1 amplitudes in response to the 140% step threshold perturbation condition were greater than in N1 amplitudes in response to 60% step threshold perturbations (p<0.001), and there was no significant interaction effect between perturbation magnitude and cohort (p = 0.125) (Fig. 2B).

**Figure 2.**
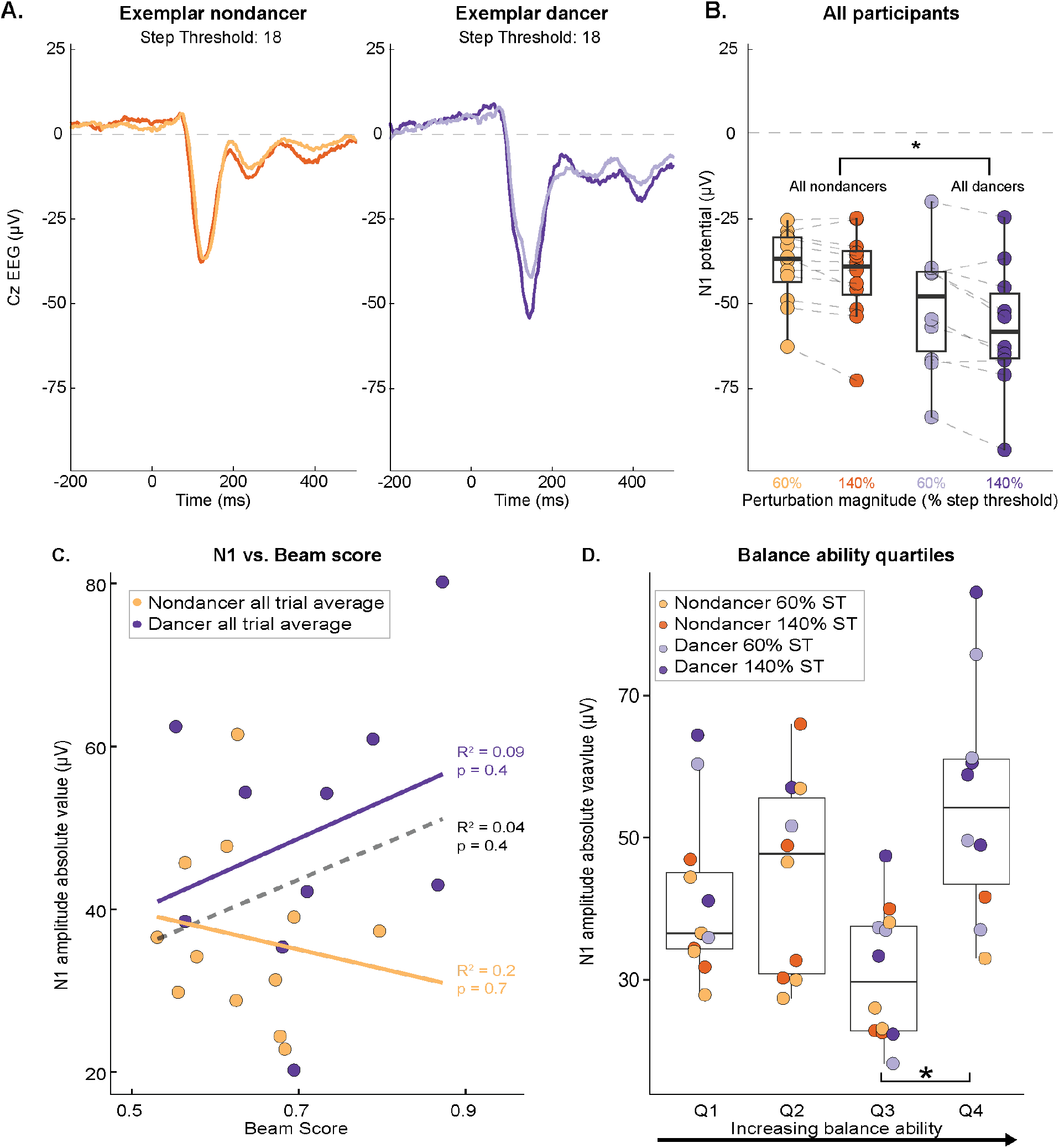
Condition average Cz ERP and N1 potential and balance ability comparison A) Condition average 60% step threshold (ST) and 140% ST ERP at the Cz electrode for an exemplar nondancer (orange) and dancer (purple) with the same ST, darker shade represents 140% condition. Perturbation onset at time = 0 ms indicated by dotted line. B) Condition average N1 potential for all participants. Asterisk indicates significant difference in potential (p < 0.05) based on a repeated measures ANOVA of N1 potential by perturbation condition and cohort. C) Individual all trial average N1 amplitude against beam score for nondancers only (orange), dancers only (purple), and all participants (black dashed line). A linear mixed effect model showed no significant effect of beam score on N1 in either cohort or condition. Linear models for each cohort individually showed no significant effect of beam score on N1 but trend positively in dancers (purple) and negatively in nondancers (orange). D) Individual condition average N1 amplitude against balance ability quartile. Points from the same participant are aligned vertically. Splitting the cohort into quartiles of balance ability, there was a significant difference between N1 amplitude in Q3 compared to Q4, which are the highest two balance ability quartiles and comprise most of the dancer cohort (n = 7 of 10 dancers). Individuals in Q4 of balance ability had larger N1 amplitudes than individuals in Q3 (*p* < 0.05).

N1 amplitude during perturbations were not correlated to balance ability, but were nevertheless higher in individuals with high balance ability (Fig. 2D). N1 amplitude was not associated with balance ability in either cohort analyzed separately (dancers p=0.4 R^2^=0.09, nondancers p=0.7,R^2^=0.2), nor together (p=0.4,R^2^=0.04) (Fig. 2C). However, when separating all participants into quartiles of balance ability rather than cohort, N1 amplitudes of individuals in the 3^rd^ and 4^th^ quartiles of balance ability were significantly different from each other (p<0.001, Fig. 2D). Within participant scaling of N1 did not differ significantly between balance ability quartiles.

We found associations between modulation of N1 responses between 60% and 140% step threshold perturbation (ΔN1) and balance ability. However, ΔN1 scaled with individual differences in step threshold in dancers (Fig. 3A), but individual differences in N1 amplitude in nondancers (Fig. 3B). Step threshold determines individual differences in the perturbation magnitudes delivered in the two conditions. Since perturbation magnitudes were a percentage of step threshold, individuals with higher step thresholds experienced an objectively larger difference between perturbation magnitudes in the 60% vs. 140% conditions. Dancers with higher step thresholds demonstrated significantly greater modulation in N1 between perturbation conditions (p=0.04) while there was no relationship between ΔN1 and step threshold in nondancers (Fig. 3A). We also investigated relationships between N1 amplitude in the 60% ST condition and ΔN1 to probe modulations in N1 proportional to subjective balance challenge (Fig. 3B). There was an association between greater modulation in N1 (ΔN1) and larger 60% ST N1 amplitude in nondancers only (p=0.08), not observed in dancers.

**Figure 3.**
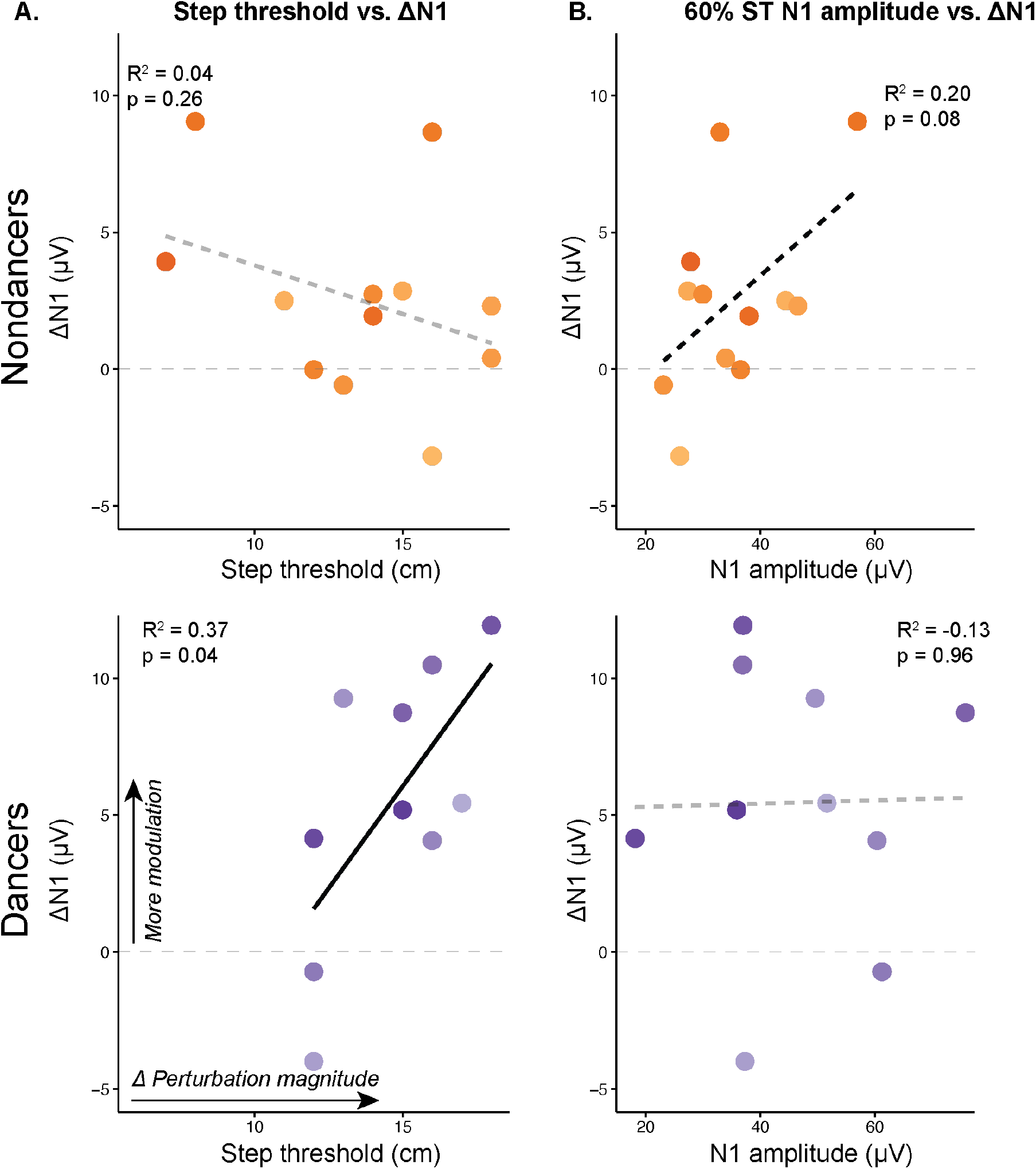
N1 modulation across cohorts with respect to step threshold and N1 at low perturbation magnitude A) Comparison of the difference in N1 amplitude between 140% and 60% ST conditions (positive ΔN1 denotes larger N1 amplitude in 140% ST perturbation) and step threshold in each cohort. A linear model showed a significant positive trend between step threshold and ΔN1 in dancers only (p=0.05) B) Comparison of ΔN1 with N1 amplitude in the 60% ST perturbation condition in each cohort. A linear model showed a near significant positive trend between N1 amplitude and ΔN1 in nondancers only (p=0.08). Shades of orange (nondancer) and purple (dancer) denote individuals.

Biomechanical outcome measures of the balance-correcting perturbation response did not differ between cohorts. Average CoP displacement in nondancers compared to dancers is similar upon visual inspection (Fig. 4A). Within participants, CoP RoR in the 140% ST condition was faster than the 60% ST condition (p=0.04) with no significant effect of cohort (p=0.45) (Fig. 4B). In non-stepping perturbations, there were no cohort level differences in peak CoP displacement or latency between dancers and nondancers (Fig. 4C). 750 ms after the perturbation, or 250 ms after the end of platform movement, increased lumbar flexion was observed in the 140% step threshold condition (p=0.4), with no effect of cohort (Fig. 4D).

**Figure 4.**
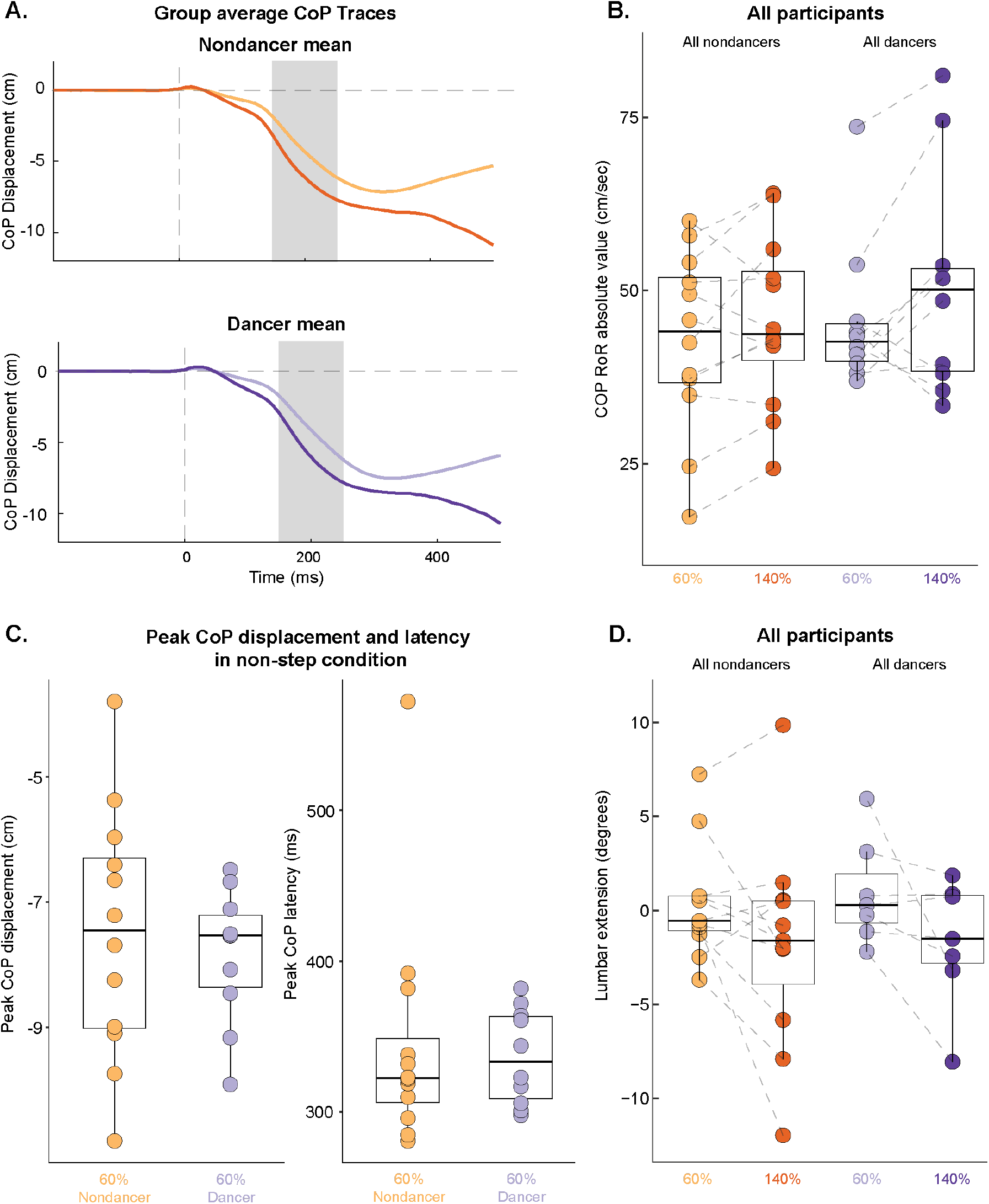
CoP and lumbar extension metrics of reactive balance response across cohorts A) Nondancer and dancer mean CoP displacement over time in response to forward perturbations at 60% step threshold (ST) and 140% ST. Perturbation onset is at time 0 ms; shaded regions indicate 150 ms and 250 ms post-perturbation, the time bin at which CoP RoR is calculated as the slope of CoP displacement for each participant and condition. B) Condition average N1 potential for all participants. A linear mixed effects model of CoP RoR by perturbation condition and cohort revealed no significant relationships between CoP and perturbation size or cohort. C) In the non-stepping condition only, peak CoP displacement and the latency of peak CoP displacement in each cohort. Paired t-tests showed no significant difference between cohorts. D) Lumbar extension angle 750 ms post-perturbation, or 250 ms after the end of platform movement. A repeated measures anova showed a significant effect of perturbation condition on lumbar extension angle (p=0.04) but not cohort.

## DISCUSSION

Our results provide evidence supporting greater cortical response sensitivity to balance errors in dancers compared to nondancers. In contrast with previous findings, individuals with the highest balance ability demonstrated the greatest cortical evoked responses (N1) to balance perturbations, which may reflect greater attention to posture and balance due to dance training. Scaling perturbations to individual balance challenge revealed a dissociation of N1 modulation, suggesting a scaling to individual task difficulty in nondancers and objective task difficulty in dancers. Despite differences in cortical engagement, balance correcting biomechanical responses to perturbations were similar across groups, highlighting the need to investigate cortical dynamics in reactive balance in addition to biomechanics in the context of both training and rehabilitation.

Modern dancers have received specific training which may lead to more sensitive assessment of balance disturbances. Due to prior results showing smaller cortical evoked potentials in individuals with better balance (33), we predicted modern dancers—a cohort of balance experts (8)—would also have smaller cortical evoked potentials in response to balance perturbations. However, the finding of larger N1 amplitudes in dancers is consistent with interpretations of N1 as an error assessment signal (39). Recent work has shown increased N1-like cortical potentials when young adult participants stood at the edge of an elevated platform, where the perceived consequence of losing balance is greater (59). In dancers, the consequence of losing balance while maintaining difficult postures, for example during a performance, is also greater than during typical standing balance. Moreover, the need to maintain a specific postural alignment is consistent with cortical response modulation in response to objective differences in perturbation magnitude. Maintaining a specific postural configuration requires alignment to external references such as the vertical, or perhaps to the postures of other dancers. In contrast, balance control requires only that the center of mass be maintained within an individual’s base of support in a gravity field, without necessitating a return to the initial posture following a perturbation (60, 61). Therefore, increased cortical responses reflecting balance error assessment may be an effect of dance training, where heightened sensitivity to balance perturbation is useful for detecting and correcting for small changes in posture (8). Whether the N1 shapes later motor output is still unclear however, so the extent to which cortical error assessment reflects overall cortical control or automaticity of balance recovery is unknown.

Cortical N1 responses may be an indicator of the degree of attention that individuals place on their balance control, regardless of their level of skill or motor impairment. In older adults, larger N1 amplitudes have been observed in individuals with a history of falls compared to those without (25), leading to the possibility of the N1 as a biomarker for fall risk (62). In individuals with a history of falls, cortical resources may serve to compensate for diminished subcortical resources (20, 37, 38, 40, 63). However, our results are consistent with the interpretation that increased cortical error assessment is not always associated with worse task performance (36). We show here that dancers may have larger evoked cortical responses, potentially enabling more precision in balance and postural control rather than reflecting a compensation for poor balance. Accordingly, a “U-shaped function” that describes greater cortical engagement at both ends of a skill level spectrum has been proposed before in balance control (44, 64). In the context of balance dysfunction, a larger N1 may indicate more sensitive cortical error assessment, indicative of greater attention to balance perturbations in individuals at high fall risk (65) or lower balance confidence (38). In able-bodied young adults, recent work has shown evidence that actively attending to balance decreases N1 amplitude (42). In individuals who have received balance training such as through dance, attending to balance may not be a conscious process, yet sensitivity to balance perturbations may still be reflected in the cortical N1 response.

Dancers have increased postural awareness (8, 66), which may bias perception toward objective task difficulty rather than individual challenge level (44). Scaling perturbations to individual step threshold allowed us to control the task for balance challenge and observe modulation in N1 amplitude between conditions of varying difficulty. As expected, scaling perturbations to step threshold reliably elicited increases in N1 amplitude with perturbation magnitude, likely because perturbations were equally challenging across participants, regardless of individual balance ability (44). In dancers, N1 amplitude was modulated proportionally to step threshold, which also drives difference in absolute perturbation magnitudes experienced across individuals. In nondancers, individual differences in N1 magnitude, or each individual’s sensitivity to balance perturbations, were correlated to difference in N1 amplitude across the two perturbation magnitudes (ΔN1 from 140% ST to 60% ST), despite objective differences in perturbation magnitudes across individuals. In contrast, dancers did not scale N1 responses based on N1 sensitivity to balance perturbations, but rather to the objective differences in perturbation magnitude. As such, dancers with similar step thresholds but different N1 amplitudes had similar N1 modulation across perturbation, whereas nondancers with similar N1 amplitude but different step thresholds had similar N1 modulation across perturbations. We interpret these results to mean that N1 responses reflect individual task difficulty in nondancers, but objective task difficulty in dancers. Since dancers are trained to maintain specific, aesthetic postures, they must know where their body is objectively, relative to each other or an audience, rather than subjectively based on their own ability or experience. Accordingly, prior work demonstrates that dancers have a preference for staying vertical, or attending to posture, than simply maintaining balance by keeping the CoM over their base of support (66, 67).

Biomechanical outcome measures could not explain differences in cortical activity between dancers and nondancers. N1 amplitudes in response to balance perturbation were larger in dancers compared to nondancers even though the biomechanics of the balance response, demonstrated here with CoP metrics (48) and trunk motion (68), did not differ between cohorts. While the biomechanical metrics analyzed here are not exhaustive and limited because of the inherent biomechanical differences in stepping and non-stepping responses (68), our results highlight the insights cortical activity can provide that may not be observable in behavior alone. Future analyses of muscle activity throughout the balance response may be able to link brain activity to behavior, as the timing of the nervous system can be observed with electromyography, potentially more sensitively than CoP and kinematic metrics (54).

Measuring cortical responses during a task that is equally challenging across individuals can provide insight into cortical contributions to reactive balance control. While we interpret N1, the most salient and rapid neurophysiological signature of the balance response, as a measure of cortical error assessment here, it may not describe the degree to which a subsequent movement is cortically or subcortically controlled (29, 35, 40, 45). Future analysis of other EEG outcome measures, such as ERP analyses during the later motor response or analysis of oscillatory activity (54, 58, 69, 70), may provide a more complete picture of cortical engagement beyond initial error assessment. Ultimately, using EEG recordings during reactive balance may help to identify or quantify individual-specific balance challenge to guide training and rehabilitation across expertise, aging, and balance impairment (37, 48, 58).

## DATA AVAILABILITY

Data that support the findings of this study are available from the corresponding author [L.H.T.] upon reasonable request.

## GRANTS

This work was supported by the National Institutes of Health, Eunice Kennedy Shriver National Institute of Child Health and Human Development, F32HD105458 (to J.L.M.); National Science Foundation, Graduate Research Fellowship Program, 1937971 (to S.E.B.); and National Institute on Aging, R01 AG072756 (to L.H.T. and M.R.B.).

## DISCLOSURES

Authors declare no conflict of interest.

## AUTHOR CONTRIBUTIONS

Conceived and designed research: K.G.K, S.E.B., L.H.T; performed experiments: K.G.K., S.E.B.; analyzed data K.G.K., S.E.B., J.L.M., J.P.; interpreted results of experiments: K.G.K., S.E.B., J.L.M., J.P., M.R.B., L.H.T.; prepared figures: K.G.K.; drafted manuscript: K.G.K.; edited and revised manuscript: S.E.B., J.L.M., J.P., M.R.B., L.H.T.; approved final version of manuscript: K.G.K., S.E.B., J.L.M., J.P., M.R.B., L.H.T.

## Notes

### Competing Interest Statement

The authors have declared no competing interest.

